# Root-derived long-range signals activate ABA synthesis in *frd3* leaves to enhance drought resistance

**DOI:** 10.1101/2023.03.07.531606

**Authors:** Qian-Qian Liu, Jin-Qiu Xia, Jie Wu, Ping-Xia Zhao, Gui-Quan Zhang, Cheng-Bin Xiang

## Abstract

Vascular plants have evolved sophisticated mechanisms of long-distance signaling to cope with environmental stress. Reactive oxygen species (ROS) act as systemic signals in plant stress responses. However, it is not known whether ROS serve as root-to-shoot signals in the drought response. Here, we show that *ferric reductase defective3* (*frd3*) mutants exhibit enhanced drought resistance concomitant with increased *NCED3* transcript levels and ABA contents in leaves. The *frd3* mutants also have an elevated hydrogen peroxide (H_2_O_2_) level in roots and leaves compared with the wild type. Grafting experiments demonstrate that drought resistance can be conferred by the *frd3* rootstock, suggesting that long-distance signals derived in *frd3* roots trigger ABA level increases in leaves and thereby enhance drought resistance and that H_2_O_2_ is a strong candidate for long-distance signals. Furthermore, comparative transcriptome and proteomics analyses revealed that many genes and proteins involved in the abiotic stress response, ROS homeostasis, and signaling pathways were affected in the *frd3* mutant, supporting the drought resistance phenotype. Taken together, our findings suggest that *frd3* root-derived long-range signals activate ABA synthesis in leaves and enhance drought resistance, indicating possible root-to-shoot H_2_O_2_ signaling in the plant drought response.

## Introduction

As sessile organisms, plants are constantly exposed to changing environments and have evolved sophisticated mechanisms to cope with these changes. Systemic acquired acclimation (SAA) (Karpinski et al., 1999; Smirnoff and Arnaud, 2019; Zhang et al., 2021) is the adaptation process of plants to abiotic stresses and is crucial for plant survival (Karpinski et al., 1999; Mittler and Blumwald, 2015).

SAA transmits signals from the sensing site to the whole plant, which is associated with signal generation, transport, and reception. To date, signals involved in rapid system transmission include ROS, calcium, hydraulic waves, and electric signals (Choi et al., 2017; Fichman et al., 2020). ROS function as important signaling molecules to regulate plant development and stress response. For instance, ROS modulate the temporal-spatial coordination of signals in cell-to-cell communication in plants (Mittler et al., 2011; Suzuki et al., 2013). Under high light and heat stress, ROS are rapidly produced at the sensing site in plants and transported to the systemic leaves, thus adapting to the surrounding changes for the whole plant (Suzuki et al., 2013; Devireddy et al., 2018). The propagation of ROS waves is dependent on the ROS-producing enzyme RESPIRATORY BURST HOMOLOG D (RBOHD), which is directly regulated by Ca^2+^ in *Arabidopsis* (Miller et al., 2009). Ca^2+^-permeable channels can also be activated by ROS in the plasma membrane, which forms a feed- forward loop, leading to more ROS production through Ca^2+^-dependent activation of RBOHs (Pei et al., 2000; Demidchik et al., 2007; Choi et al., 2016). ROS autocatalysis is well suited to rapidly build ROS levels in different tissues through the transmission of ROS waves for signal amplification (Talaat, 2019).

Long-distance signal transduction between above- and belowground parts plays an important role in the drought response of plants. Roots are generally thought to perceive surrounding changes primarily and transmit signals to the whole plant to trigger adaptation (Li et al., 2021). ABA was suggested to act as a key stress response hormone that participates in communication from the root to shoot upon soil drying (Wilkinson and Davies, 2002). However, it has been demonstrated that ABA-induced stomatal closure does not require ABA from roots (Holbrook et al., 2002; Christmann et al., 2007; Goodger and Schachtman, 2010). Hydraulic signals have been shown to function in root-to-shoot communication to activate ABA biosynthesis in leaves under dehydration stress (Christmann et al., 2007). Recently, the CLAVATA3/EMBRYO-SURROUNDING REGION-RELATED25 (CLE25) peptide has been reported to function as a root-to-shoot signal and interacts with BAM1/3 to enhance ABA accumulation by inducing the expression of *NCED3* (McLachlan et al., 2018; Takahashi et al., 2018). Additionally, drought-enhanced xylem sap sulfate is incorporated into cysteine to trigger ABA production and ROS accumulation, thereby affecting stomatal closure (Malcheska et al., 2017; Batool et al., 2018). Drought stress triggers the rapid accumulation of ROS in the apoplast, which is critical for stomatal closure (Qi et al., 2017; Qi et al., 2018). HYDROGEN-PEROXIDE-INDUCED Ca^2+^ INCREASE (HPCA), as a sensor of H_2_O_2_ in guard cells, was shown to modulate stomatal closure (Wu et al., 2020). Moreover, as long-distance signals, ROS/Ca^2+^ waves are required for regulating ABA responses and stomatal closure by activating GUARD CELL HYDROGEN PEROXIDE-RESISTANT1 (GHR1) and the S-type anion channel SLAC1 in systemic leaves to light stress (Devireddy et al., 2018; Kollist et al., 2019; Takahashi et al., 2020). Nevertheless, long-distance ROS signaling, especially root-to-shoot signaling, in the drought response of plants has not been well addressed.

ROS play a dual role in plant growth, not only functioning as signaling molecules but also generating toxic effects in cells. Iron (Fe) acts as the most important transition metal ion catalyzing ROS generation and elimination (Le et al., 2019). It has been found that excessive accumulation of iron in the roots can cause an increase in ROS (Finatto et al., 2015; Zhang et al., 2018). Moreover, improving iron uptake can promote plant adaptation to adverse environments (Pirzad and Shokrani, 2012; Tripathi et al., 2018; Dvořák et al., 2021). Foliar Fe application with Zn diminishes oxidative stress by reducing ROS levels and lessening lipid peroxidation by enhancing antioxidant enzymes (CAT, GPX, and SOD) under drought stress (Akbari et al., 2013; Amirinejad et al., 2013). In addition, high levels of labile Fe^2+^ in cells perturbed the ROS balance by producing more hydroxyl radicals (Mittler, 2017).

Dicotyledonous plants have evolved mechanisms to uptake iron from the soil. First, Fe^3+^ is activated by the release of protons mediated by H^+^-ATPase (AHA2) (Santi and Schmidt, 2009). Mobilized Fe^3+^ is reduced to Fe^2+^ by a membrane-bound Fe^3+^ reductase oxidase (FRO2) and then transported into the root epidermal cells by IRT1 (Brumbarova et al., 2015; Curie and Mari, 2017). Several studies have demonstrated the importance of Fe uptake by chelation. Citrate acts as an iron chelator and plays an important role in iron transport from roots to shoots via xylem sap (Alvarez-Fernandez et al., 2014). Fe^3+^-citrate is the main form of iron translocated in the xylem (Bruggemann et al., 1993; Rellan-Alvarez et al., 2010; Kryvoruchko et al., 2018). Iron uptake and transport were more efficient in *Arabidopsis* with the overproduction of citrate under Fe-deficient conditions (Ramirez-Rodriguez et al., 2001). Meanwhile, citrate levels were significantly increased in pea roots under Fe deficiency (Jelali et al., 2010; Kabir et al., 2012). A recent study revealed that citrate participates in nitrate-mediated iron deficiency in apples (Sun et al., 2021).

Citrate is crucial for iron transport. Ferric reductase defective 3 (FRD3), a member of the multidrug and toxic compound extrusion (MATE) family in *Arabidopsis*, mediates the efflux of citrate from intracellular to apoplast (Rogers and Guerinot, 2002; Roschzttardtz et al., 2011; Pineau et al., 2012). The *frd3* mutants show reduced excretion of citrate and accumulate excess Fe in roots, causing shoot Fe deficiency and chlorosis (Green and Rogers, 2004; Durrett et al., 2007; Li et al., 2019).

*OsFRDL1*, an orthologous gene of *AtFRD3* in rice, is expressed in root pericycle cells and is essential for long-distance Fe translocation as an Fe-citrate complex (Inoue et al., 2004; Yokosho et al., 2009). It is not well understood whether citrate transport is involved in the drought response in plants. We previously identified a candidate gene responsible for the enhanced drought resistance of a single-segment substitution line in rice. The candidate gene was predicted to encode a citrate transporter. To facilitate the investigation of how a defective citrate transporter enhances drought resistance, we explored the functions of the *Arabidopsis* citrate transporter *FRD3* in drought resistance. In this study, we report that loss of *FRD3* leads to drought resistance in *Arabidopsis*, which was attributed to elevated *NCED3* expression and leaf ABA levels. The *frd3* mutants accumulated higher H_2_O_2_ levels in roots and leaves than the wild type. Grafting experiments showed that root-to-shoot signals, likely ROS resulting from *FRD3* disruption, activated ABA synthesis in leaves, leading to a drought-resistant phenotype. Furthermore, transcriptome and proteomics data support that loss of *FRD3* alters the expression levels of the genes and proteins involved in the drought stress response and ROS homeostasis, which contribute to drought tolerance. Our findings indicate that root-derived long-range signals can trigger a drought response cascade in the leaves of *frd3* mutants, providing new insights into the long-distance signal-mediated drought response in plants.

## Results

### Loss of *FRD3* leads to improved drought resistance

To investigate the function of *FRD3* in drought resistance, we obtained a T-DNA insertion mutant *frd3-8* (SALK_122235) and the mutant *man1-1* (CS8506) with a single recessive mutation in *FRD3* from ABRC and constructed *FRD3* overexpression lines OX-3 and OX-4. These lines were confirmed before the soil drought resistance assay (Figure S1). Under normal conditions, all the genotypes displayed similar growth and development as the wild type except that loss-of-*FRD3* mutants showed leaf chlorosis as previously reported (Roschzttardtz et al., 2011; Charlier et al., 2015; Scheepers et al., 2020). However, after drought treatment for 15 days, OX-3, OX-4, and the wild-type plants severely wilted and exhibited a survival rate of approximately 30%, whereas *frd3-8* and *man1-1* mutants only showed slight wilting and had a survival rate of approximately 90% (Figure 1A and B). Moreover, the water loss from the soil potted with OX lines and wild type was similar but significantly faster than that with the mutants (Figure 1C). These results indicate that the loss of *FRD3* enhances drought resistance.

**Figure 1.**
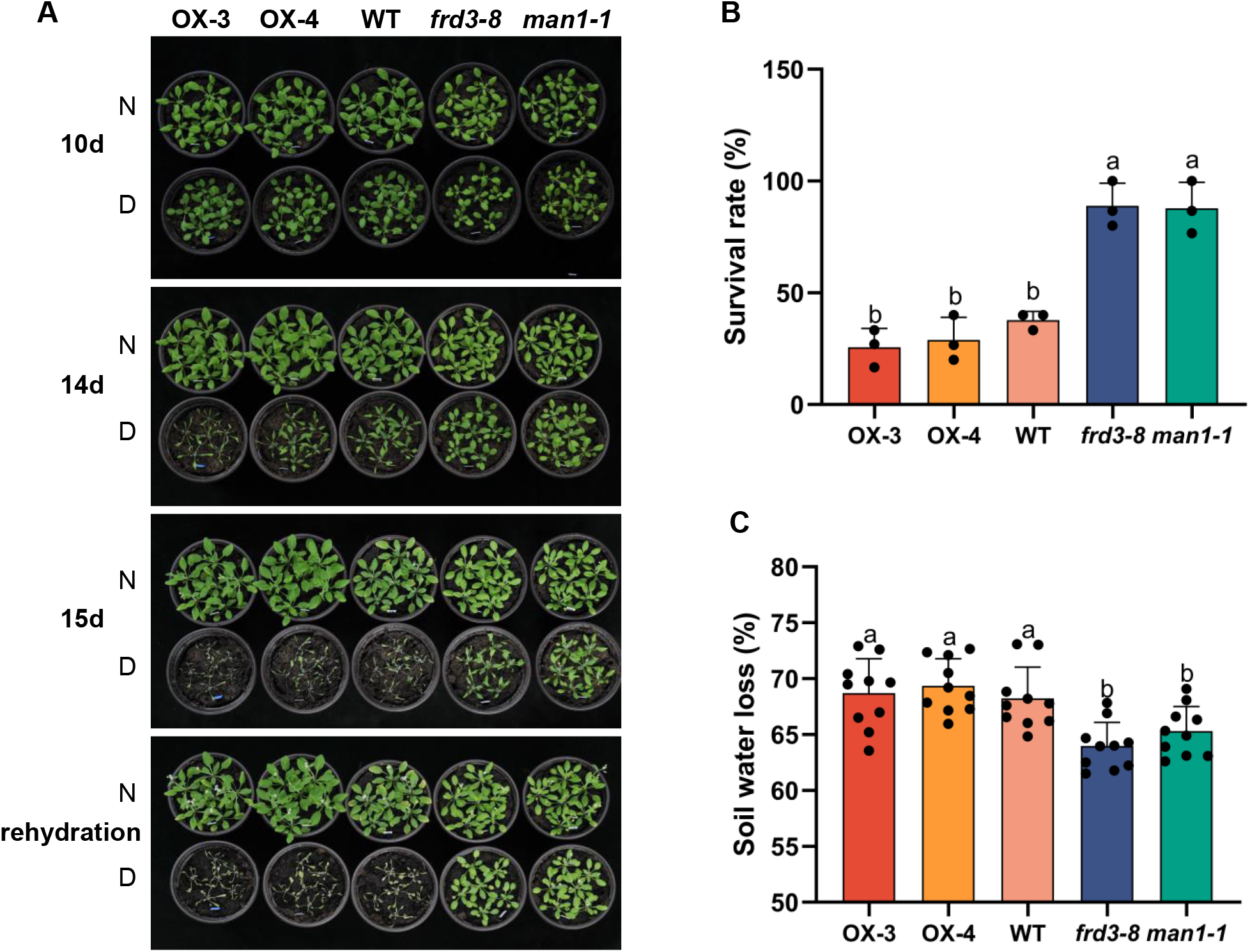
Loss of *FRD3* enhances drought resistance. A-B. Drought resistance assays. 1-week-old wild type (WT), overexpression lines (OX-3 and OX-4), and the knockout mutants *(frd3-8* and *man1-1*) plants grown under normal conditions received drought treatment (without watering, D) or normal irrigation (N) for 15 days before rehydration. Photographs were taken respectively at 10, 14, and 15 days, and after rehydration (A). The survival rate was calculated after rehydration (B). Values are mean ±SD (n ≥ 3 replicates, 60 seedlings per replicate). Different letters indicate significant differences by one-way ANOVA (*P* < 0.05). C. Soil water loss. The initial weight (m_i_) and the weight after 15 days of drought (m_15d_) were recorded and the soil water loss rate ((m_i_ - m_15d_) / m_i_) was calculated. Values are mean ±SD (n ≥ 3 replicates, 10 pots per genotype). Different letters indicate significant differences by one-way ANOVA (*P* < 0.05).

### Loss of *FRD3* alters stomatal density and ABA content in leaves

To explore the mechanism by which *FRD3* contributes to drought resistance, we examined stomatal density, stomatal index, and stomatal aperture. Compared with the wild type, *frd3-8* and *man1-1* mutants exhibited a lower stomatal density and stomatal index, whereas the overexpression line OX-3 showed similar stomatal density and index as the wild type (Figure 2A-C). We also examined the sensitivity of stomatal movement to ABA treatment. Under normal conditions, there was no significant difference in stomatal aperture between the wild-type, *frd3-8*, *man1-1*, and OX-3 lines. Nonetheless, in response to 5 µM ABA treatment, the stomatal aperture of *frd3* mutants was significantly smaller than that of the wild type and overexpression line (Figure 2D), suggesting that the guard cells of the mutants are more sensitive to ABA.

**Figure 2.**
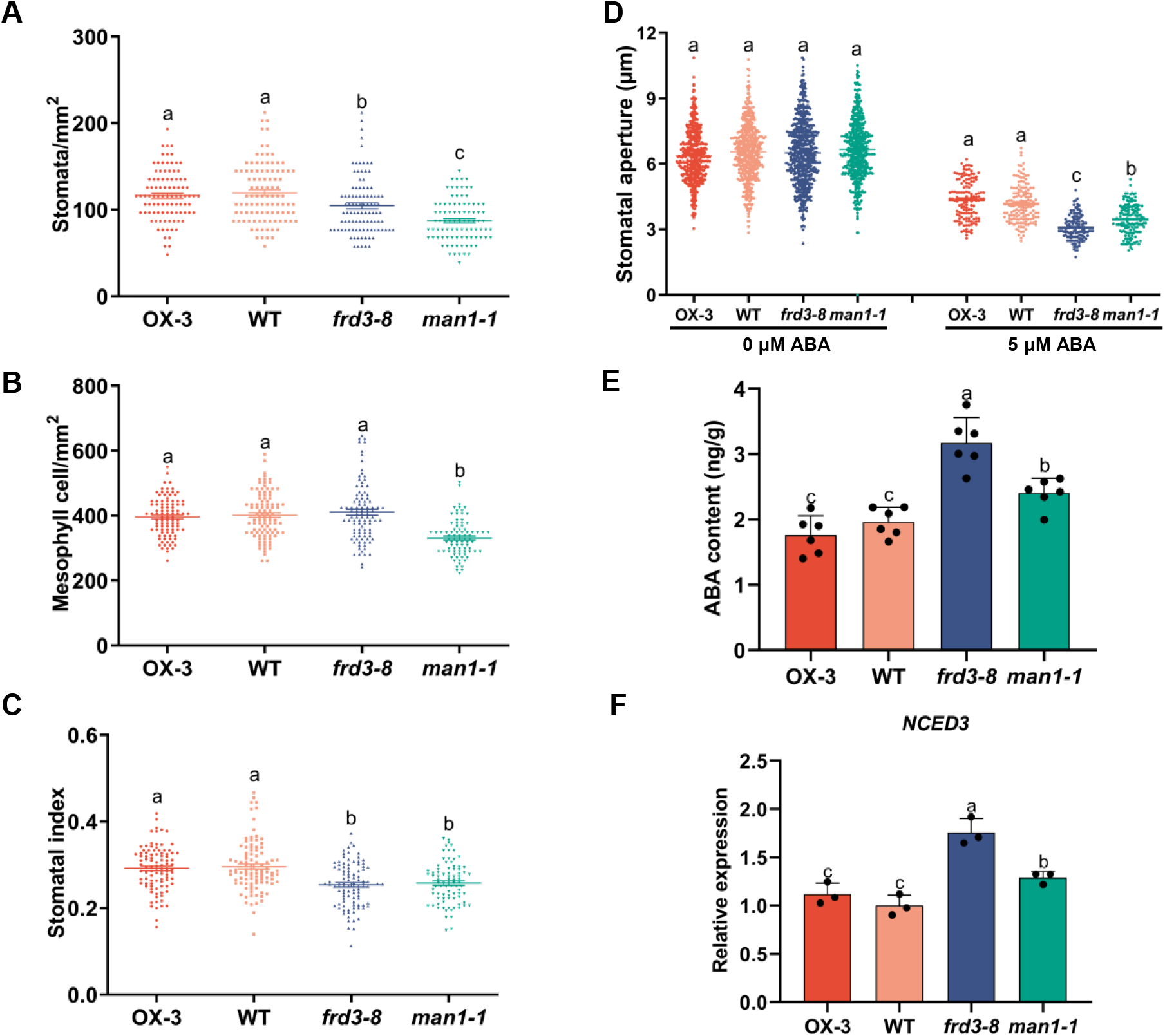
Loss of *FRD3* alters stomatal density and ABA content in the leaves. A-D. Under normal conditions, The abaxial epidermis of 3-week-old soil-grown plants of OX-3, WT, *frd3-8*, and *man1-1* lines were obtained. The number of stomata and mesophyll cells was recorded under the microscope, and the stomatal density (A), mesophyll cells density (B), and stomatal index (the ratio of stomata to epidermal cells plus stomata, C) was calculated, respectively. Stomatal aperture was measured using ImageJ software (D). Values are mean ±SD (n ≥ 3 replicates, 10 seedlings per genotype, two leaves were taken from each plant, and at least five fields of view were taken from each leaf). Different letters indicate significant differences by one-way ANOVA (*P* < 0.05). E. leaves ABA content of 3-week-old soil-grown OX-3, WT, *frd3-8*, and *man1-1* lines under normal conditions was checked using ELISA. Values are means ±SD (n ≥ 3 replicates, 10 seedlings per genotype). Different letters indicate significant differences by one-way ANOVA (*P* < 0.05). F. The transcript levels of *NCED3* in the leaves of 3-week-old soil-grown OX-3, WT, *frd3-8*, and *man1-1* lines under normal conditions were measured by qRT‒PCR. Values are mean ±SD (n ≥ 3 replicates). Different letters indicate significant differences by one-way ANOVA (*P* < 0.05).

To further verify whether the ABA level is altered in different *FRD3* genotypes, we measured the ABA content in wild-type, *frd3-8*, *man1-1*, and OX-3 lines. The ABA content in *frd3-8* and *man1-1* mutants was significantly higher than that in the wild type under normal conditions, while OX-3 had a similar ABA content as the wild type (Figure 2E). Additionally, we found that the expression of *NCED3*, which is involved in ABA biosynthesis, was significantly elevated in *frd3-8* and *man1-1* (Figure 2F). Together, these results indicate that the loss of *FRD3* increases leaf ABA levels, reduces stomatal density, and increases the sensitivity of guard cells, thereby improving drought resistance.

### Loss of *FRD3* increases H_2_O_2_ accumulation

The loss of *FRD3* is known to cause ROS accumulation in the roots (Scheepers et al., 2020). To further explore the drought resistance mechanism of the *frd3* mutants, we examined the ROS levels in the wild-type, *frd3-8*, *man1-1*, and OX-3 lines by diaminobenzidine (DAB) and nitro blue tetrazolium (NBT) staining. The DAB staining in the roots of *frd3-8* and *man1-1* mutants was much darker compared with that of the wild type (Figure 3A, top). Similar results were observed in rosette leaves (Figure 3A, bottom). The relative staining intensity in the roots and leaves of *frd3* mutants was significantly higher than that of the wild type and OX-3 lines (Figure 3B). Nevertheless, no obvious difference was observed in NBT staining in roots or leaves among the genotypes (Figure 3C and D). Taken together, these results show that loss of *FRD3* causes higher H_2_O_2_ accumulation in roots and leaves, suggesting that H_2_O_2_ might act as a long-distance signal that is derived in roots and translocated to leaves, where it triggers an increase in ABA levels and eventually leads to the drought resistance phenotype.

**Figure 3.**
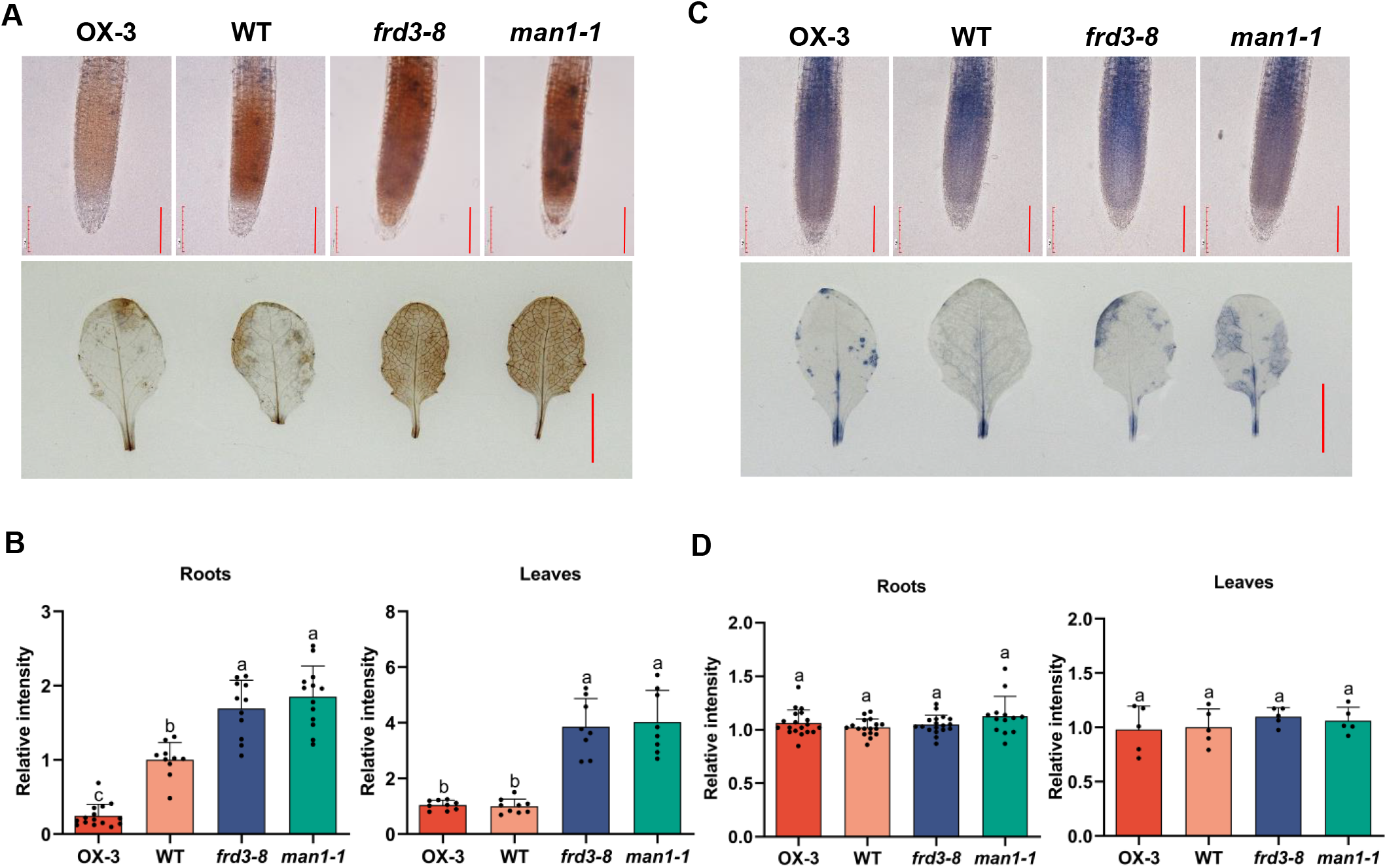
Loss of *FRD3* increases hydrogen peroxide accumulation in the roots and leaves. A-B. Hydrogen peroxide accumulation in OX-3, WT, *frd3-8*, and *man1-1* lines. One-week-old seedlings grown on MS medium and 3-week-old detached rosette leaves grown on soil were incubated in DAB (A) solution and stained for 3 hours (seedlings) or overnight (rosette leaves) in darkness, then de-staining before photographs of roots tips and leaves were taken. Bar = 100 μm in roots, bar = 1 cm in leaves. The relative intensity in the roots and leaves of different lines was counted using ImageJ software (B). The experiment was repeated at least three times (≥10 seedlings per replicate). Different letters indicate significant differences by one-way ANOVA (*P* < 0.05). C-D. NBT staining. One-week-old seedlings grown on MS medium and 3-week-old detached rosette leaves grown on soil were incubated in NBT (C) solution and stained for 0.5 hours (seedlings) or overnight (rosette leaves) in darkness, then de-staining before photographs of roots tips and leaves were taken. Bar = 100 μm in roots, Bar = 1 cm in leaves. The relative intensity in the roots and leaves of different lines was counted using ImageJ software (D). The experiment was repeated at least three times (≥10 seedlings per replicate). Different letters indicate significant differences by one-way ANOVA (*P* < 0.05).

### Grafting experiments demonstrate that root-derived signals activate ABA synthesis in leaves and enhance drought resistance in the *frd3* mutant

To explore the roles of *FRD3* in drought resistance, we examined the expression patterns of *FRD3*. The results of qRT PCR and GUS reporter analyses revealed that *FRD3* was mainly expressed in the stele of roots and hypocotyl, where the junction between the above-ground and roots was also located (Figure S2A and B). This result suggests that FRD3, a MATE transporter exporting citrate, is likely involved in the increase in H_2_O_2_ levels in the roots and leaves of the mutants through iron transport.

To test this hypothesis, we performed grafting experiments to verify whether the root-derived signals from the *frd3* mutant rootstock could confer drought resistance to the wild-type scion. Interestingly, the *frd3* rootstock conferred similar drought resistance as the *frd3* mutant to all scions regardless of their genotypes, whereas the wild-type rootstock did not (Figure 4A and B). Moreover, the *frd3* rootstock also increased ABA content, *NCED3* transcript level, and H_2_O_2_ level in the leaves of the wild-type scion, as in the *frd3* mutant (Figure 4C-E). Taken together, these grafting experiment results demonstrate that *frd3* root-derived signals trigger a series of events in the leaves of grafted scions, including an increase in H_2_O_2_ levels, the upregulation of *NCED3*, an increase in ABA content, and ultimately enhanced drought resistance. The most likely candidates for root-derived signals include elevated H_2_O_2_ and Fe deficiency.

**Figure 4.**
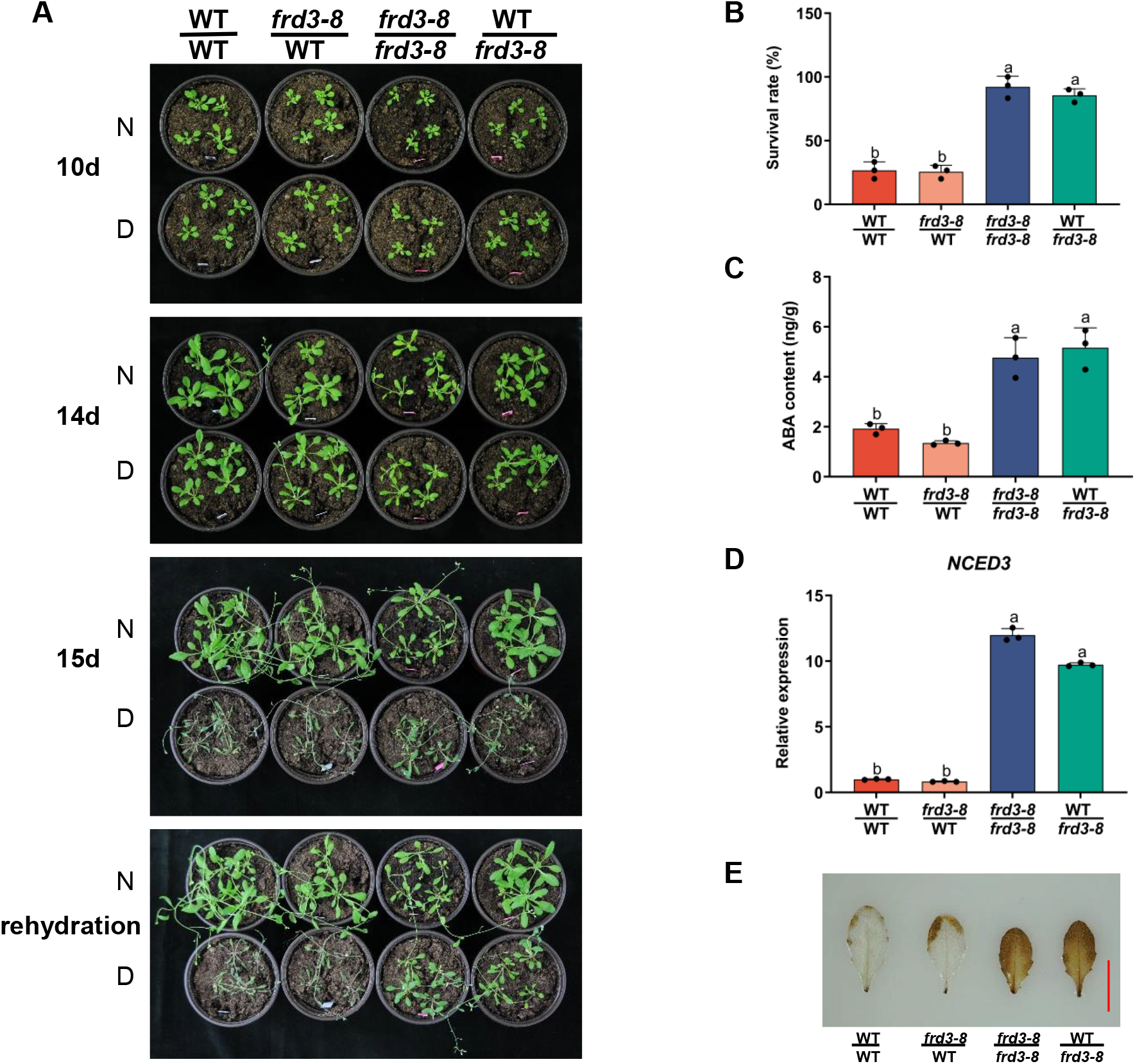
Root-derived signals activate ABA synthesis in the leaves and enhance drought resistance. A-B. Drought resistance assays of grafted plants. The grafted plants, as indicated by scion/rootstock genotypes, were grown in soil for 10 days (normal irrigation) and then subjected to drought treatment (without watering, D) or normal irrigation (N) for 15 days before rehydration. Photographs were taken respectively at 10, 14, 15 days, and rehydration (A). The survival rate was counted one day after rehydration (B). Values are mean ± SD (n ≥ 3 replicates, 30 seedlings per genotype). Different letters indicate significant differences by one-way ANOVA (*P* < 0.05). C. Leaves ABA content of 3-week-old soil-grown grafted seedlings under controlled conditions was measured using ELISA. Values are mean ± SD (n ≥ 3 replicates, 10 seedlings per genotype). Different letters indicate significant differences by one-way ANOVA (*P* < 0.05). D. The transcript level of *NCED3* in the leaves of 3-week-old soil-grown grafted plants under normal conditions was measured by qRT‒PCR. Values are mean ± SD (n ≥ 3 replicates, 10 seedlings were collected as a replicate). Different letters indicate significant differences by one-way ANOVA (*P* < 0.05). E. DAB staining of grafted seedlings. The leaves of 3-week-old soil-grown grafted plants were incubated in DAB staining for overnight in darkness. The leaves were decolorized at least three times using decolorizing solution, then photographs were taken. The experiment was repeated at least three times (≥ 10 seedlings per replicate).

### Foliar Fe spraying cannot reverse the *frd3* drought resistance phenotype to the wild type

Fe deficiency in the leaf is typical of *frd3* mutants and could be a cause leading to the drought resistance phenotype. To rule out this possibility, we carried out a foliar Fe supplementation experiment. Nevertheless, foliar Fe supplementation failed to restore the *frd3-8* drought resistance phenotype to that of the wild type (Figure 5A and B), whereas foliar Fe application restored the leaf chlorosis and chlorophyll content of the *frd3-8* mutant to wild-type levels (Figure 5C), indicating that foliar Fe application was effective. These results suggest that leaf Fe deficiency in the *frd3* mutant is not the cause of drought resistance and reaffirm that root-derived long-range signals are responsible for drought resistance.

**Figure 5.**
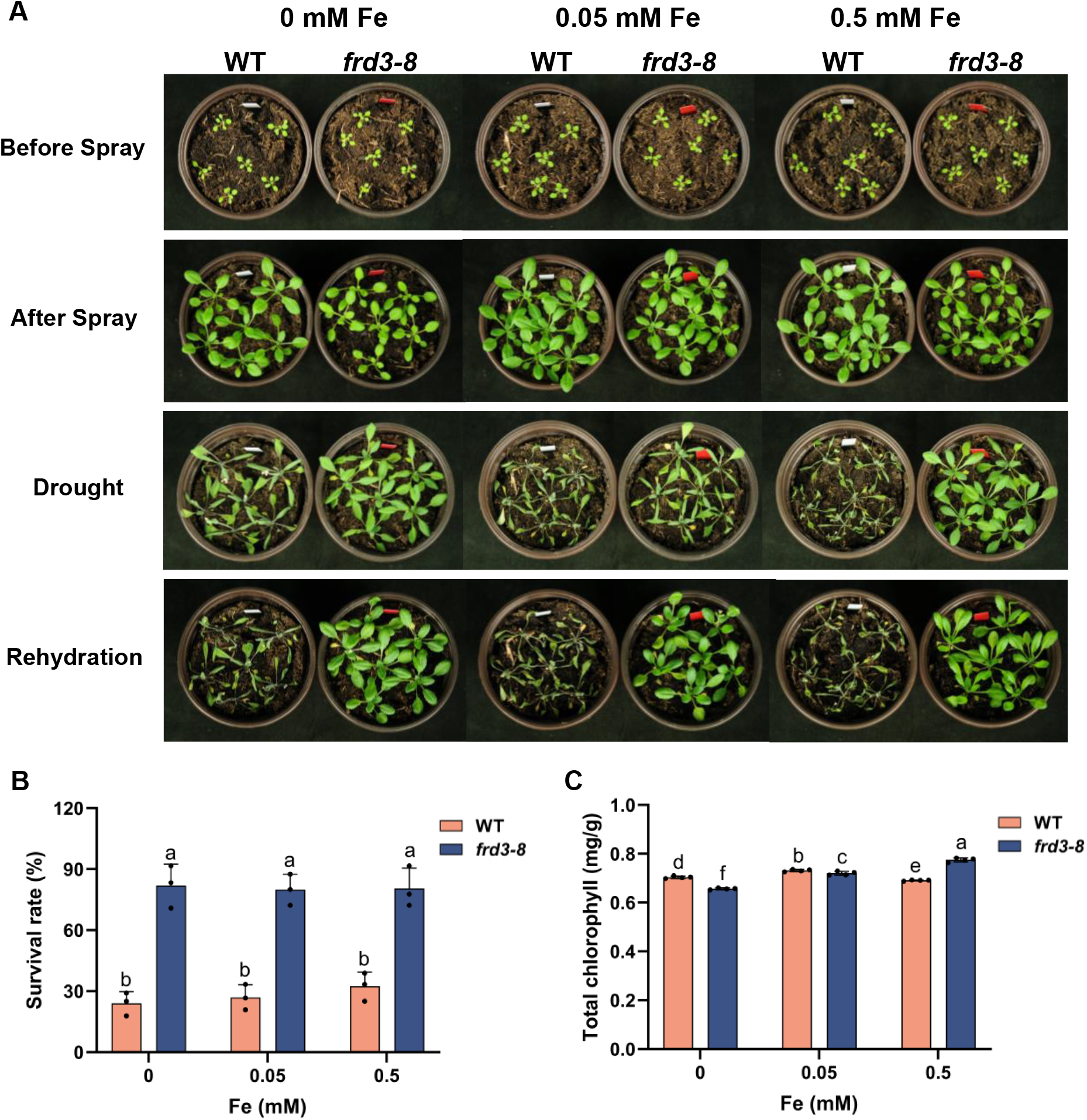
Foliar Fe supplement rules out Fe or Fe-originated signals to enhance drought resistance in *frd3-8*. A-C. Phenotype analysis of drought after foliar Fe supplement. Seedlings grown in the soil for 7 days were sprayed with 0.15 mL solutions containing 0, 0.05, and 0.5 mM Fe every 36h, 7 times in total, and then the leaf chlorophyll content was measured (C). Subsequently, drought experiments were conducted. The photograph was taken (A) and the survival rate (B) was analyzed after rehydration. The experiment was repeated at least three times (≥ 15 seedlings per replicate). Different letters indicate significant differences by one-way ANOVA (*P* < 0.05).

### Dehydration triggers H_2_O_2_ accumulation in wild-type roots and leaves

The above results suggest that root-derived signals, H_2_O_2_ being a strong candidate, trigger a series of events leading to a drought resistance phenotype in *frd3* mutants. We asked whether the wild type could generate such signals when plants experience dehydration. To probe this possibility, we dehydrated the wild-type seedlings in the air for 5 to 15 minutes as described in the methods. In contrast to the well-hydrated conditions, dehydration treatment increased H_2_O_2_ levels in roots and leaves, as shown by DAB staining (Figure 6A and B). The relative staining intensity is shown in Figure 6C and 6D. These results show that dehydration can increase H_2_O_2_ levels in the root and leaves, suggesting that H_2_O_2_ may serve as a long-distance signal in the drought response.

**Figure 6.**
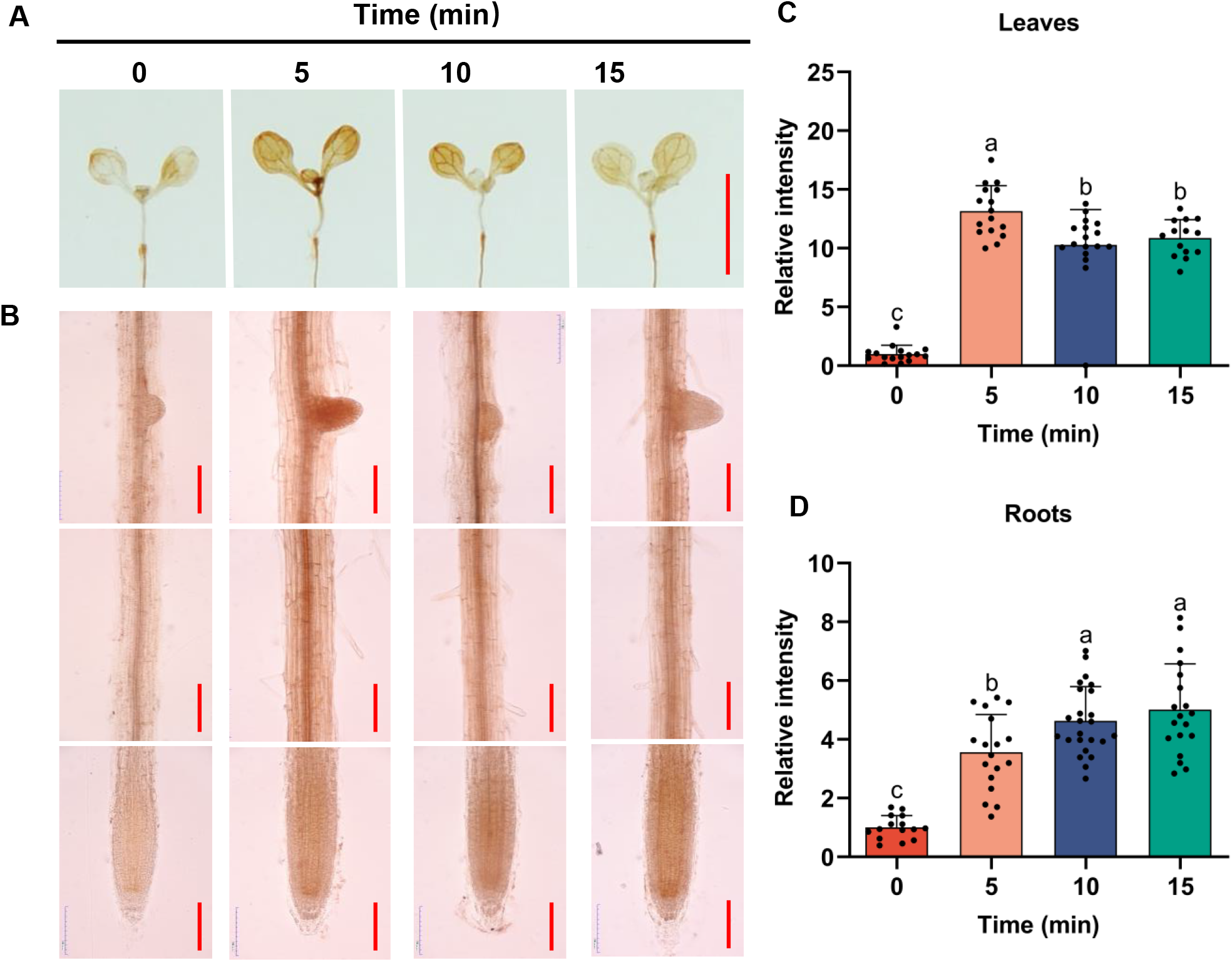
Dehydrated generates H_2_O_2_ accumulation in the wild type roots and leaves. A-B. H_2_O_2_ accumulation in the wild-type leaves and roots. Wild-type seedlings grown in MS medium for 7 days were exposed to air drying for 0, 5, 10, 15 min, and then seedlings were incubated in DAB solution for 0.5 h (roots) or 3 h (leaves) in darkness before taken photographs. Bar = 0.5 cm in A, bar = 200 μm in B. C-D. The relative staining intensity in the leaves (C) and roots (D) of wild-type was counted using ImageJ. The experiment was repeated at least three times (≥ 15 seedlings per replicate). Different letters indicate significant differences by one-way ANOVA (*P* < 0.05).

### Transcriptome analyses reveal altered expression profiles between grafted plants

To gain insights into the molecular mechanism by which loss of *FRD3* enhances drought tolerance, we performed transcriptome analyses to profile the differences in transcripts in the shoots of grafted WT/WT, *frd3-8*/*frd3-8*, WT/*frd3-8* and *frd3-8*/WT plants. The number of differentially expressed genes (DEGs) differed significantly between WT/WT vs *frd3-8*/*frd3-8* and WT/WT vs WT/*frd3-8*, but a few DEGs were identified between WT/WT vs *frd3-8*/WT and *frd3-8*/*frd3-8* vs WT/*frd3-8* (Figure 7A), suggesting that DEG expression was strongly affected in plants with *frd3* mutant rootstock. Comparative analysis of the DEGs between the WT/WT vs *frd3-8*/*frd3-8*, WT/WT vs *frd3-8*/WT, and WT/WT vs WT/*frd3-8* groups revealed only a small number of clustered genes (Figure 7B). These data demonstrated that the *frd3* rootstock widely affects the transcript profiles.

**Figure 7.**
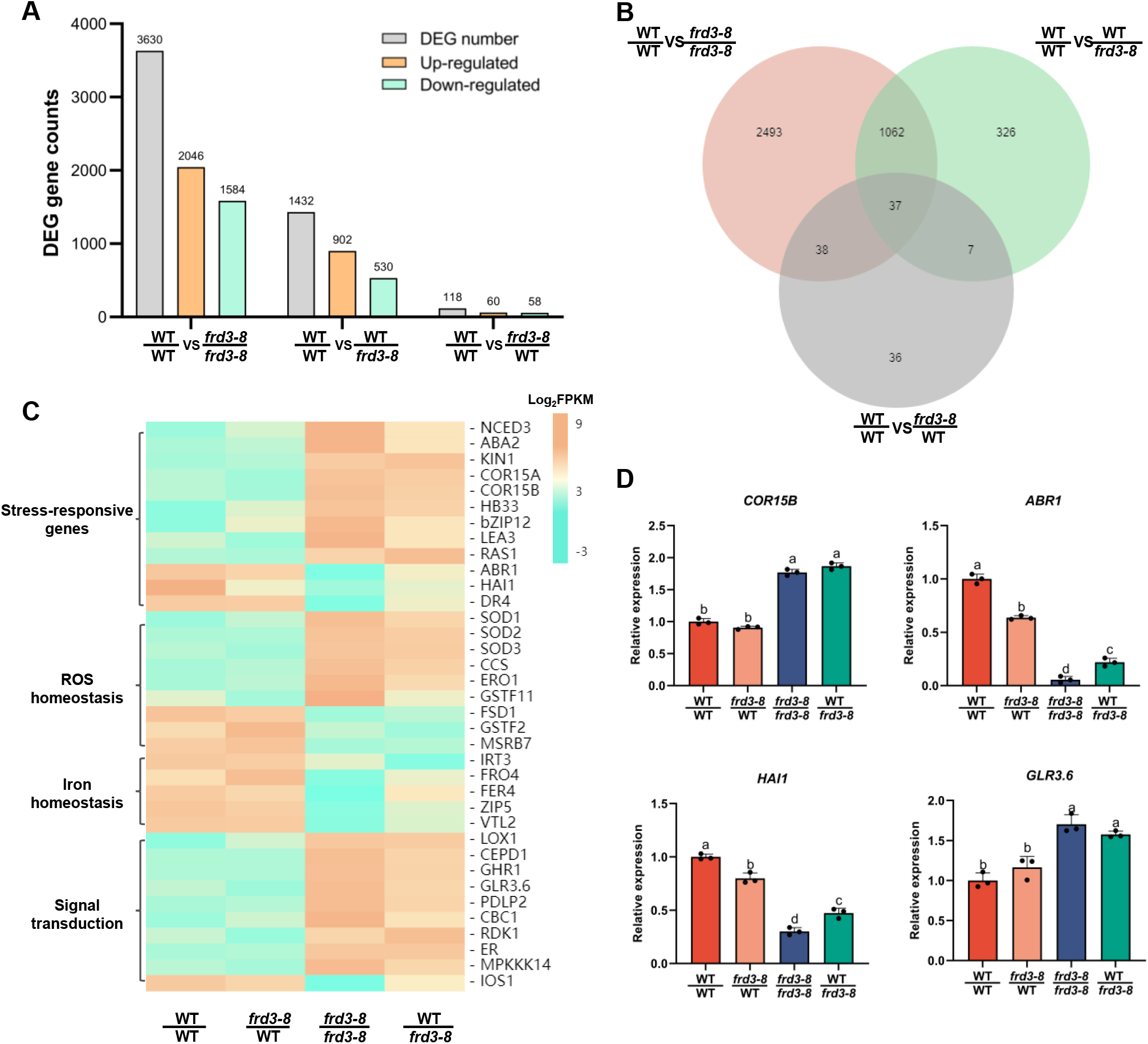
DEGs affected in different grafted lines were identified by transcriptomic analyses. A. The number of DEGs. The statistics data of DEGs in WT/WT, *frd3-8*/*frd3-8*, WT/*frd3-8,* and *frd3-8*/WT grafted plants. B. Venn diagrams analysis of the common and specific DEGs between (WT/WT vs *frd3-8*/*frd3-8*), (WT/WT vs *frd3-8* /WT), and (WT/WT vs WT/*frd3-8*) groups. The numbers represent the total numbers of DEGs in different comparison groups. C. Hierarchical clustering analysis of the key DEGs in WT/WT, *frd3-8* /WT, *frd3-8*/*frd3-8*, and WT/*frd3-8* grafted plants. The results were presented as log_2_FPKM with a *P* value < 0.05 and were confirmed with three independent experiments. FPKM: Fragments Per Kilobase of transcript per Million fragments mapped. D. The transcript levels of *COR15B*, *ABR1*, *HAI1,* and *GLR3.6* were measured in the leaves of different grafted plants by qRT‒PCR. Values are mean ±SD (n = 3 replicates). Different letters indicate significant differences by one-way ANOVA (*P* < 0.05).

To identify genes involved in signaling and metabolic pathways, we analyzed the DEGs based on the KEGG pathway in depth. Several categories, including ribosome, photosynthesis, plant hormone signal transduction, and DNA replication, were enriched in the WT/WT vs *frd3-8*/*frd3-8* and WT/WT vs WT/*frd3-8* groups (Figure S3). These pathways play an important role in the growth and development of plants and the response to stress.

To gain a global view of loss-of-*FRD3-*affected gene expression, we analyzed the expression profile of DEGs in different grafted plants by hierarchical clustering. As shown in Figure 7C, the key genes related to ABA biosynthesis, including *NCED3* and *ABA2*, and abiotic stress-responsive genes, such as *KIN1*, *COR15A*, and *COR15B*, were upregulated in the *frd3-8*/*frd3-8* and WT/*frd3-8* plants. Meanwhile, those genes involved in response to ABA, including *HB33*, *bZIP12*, *LEA3*, and *RAS1*, were higher in the *frd3-8*/*frd3-8* and WT/*frd3-8* plants, while *HAI1*, a negative regulator of ABA signaling, was downregulated. Remarkably, the transcript levels of genes encoding superoxide dismutase that function in antioxidation, including *SOD1*, *SOD2*, *SOD3*, and superoxide dismutase copper chaperone *CCS*, were significantly higher in the *frd3-8*/*frd3-8* and WT/*frd3-8* plants but lower in the WT/WT and *frd3-8*/WT plants.

The expression levels of *IRT3*, *FRO4*, and *FER4*, which are involved in iron homeostasis, were strongly inhibited in *frd3-8*/*frd3-8* plants and decreased in WT/*frd3-8* plants. In addition, a large number of genes associated with signal transduction, including *GHR1*, *GLR3.6*, *RDK1*, *ER*, and *MPKKK14,* were affected in the plants with *frd3* mutant rootstock.

To confirm the RNA-seq data, we checked the expression profiles of several genes involved in abiotic stress response and signal transduction in WT/WT, *frd3-8*/WT, *frd3-8*/*frd3-8*, and WT/*frd3-8* by qRT PCR. The transcript levels of *COR15B*, *ABR1*, *HAI1,* and *GLR3.6* were in agreement with the RNA-seq results (Figure 7D). These data suggest that loss of *FRD3* causes drought resistance by affecting the stress response, ROS homeostasis, and signal transduction. Notably, the *frd3* mutant rootstock confers a similar expression profile in the wild-type scion as the *frd3* mutant, reaffirming root-derived signals for drought resistance of *frd3*.

### Proteomics analyses show that FRD3 alters the abundance of proteins associated with the response to ROS

We performed proteomics analyses to identify the differentially expressed proteins between the *frd3-8* mutant and wild type. By comparative analysis, 148 upregulated proteins and 125 downregulated proteins were identified (Figure 8A and B). Functional classification showed that differentially expressed proteins were mostly located in the chloroplast (Figure S4), which is consistent with the chlorosis of the *frd3-8* mutant. Furthermore, GO analysis based on biological processes revealed that several categories of upregulated proteins were significantly enriched in the WT vs *frd3-8* group, including those in the response to calcium ion, vascular transport, phloem transport, and response to superoxide (Figure 8C). Additionally, the downregulated proteins were associated with cellular oxidant detoxification, ion homeostasis, response to hydrogen peroxide, and glycoside biosynthetic process (Figure 8D). Among those proteins, copper/zinc superoxide dismutase (SOD1, SOD2) and CCS, which are associated with the scavenging of superoxide radicals, were significantly increased in the *frd3-8* mutant (Figure 8E), consistent with the transcriptomic data (Figure 7C). Moreover, FER1, FER3, and FER4, which are involved in Fe homeostasis, were downregulated in the *frd3-8* mutant according to hierarchical clustering analysis. We also checked the transcript levels of *SOD2* and *CCS* in the WT, *frd3-8*, and *man1-1* lines by qRT PCR analysis. As shown in Figure 8F, *SOD2* and *CCS* were upregulated in *frd3-8* and *man1-1* mutants, consistent with the proteomics analysis. Additionally, SOD enzyme activity in *frd3-8* and *man1-1* mutants was higher than that in WT (Figure 8G). Taken together, these results show that loss of *FRD3* alters the levels of proteins associated with vascular/phloem transport and response to superoxide/hydrogen peroxide In particular, proteins associated with ROS scavenging were upregulated in the *frd3*-8 mutant, which is consistent with the transcriptomic results (Figure 7C).

**Figure 8.**
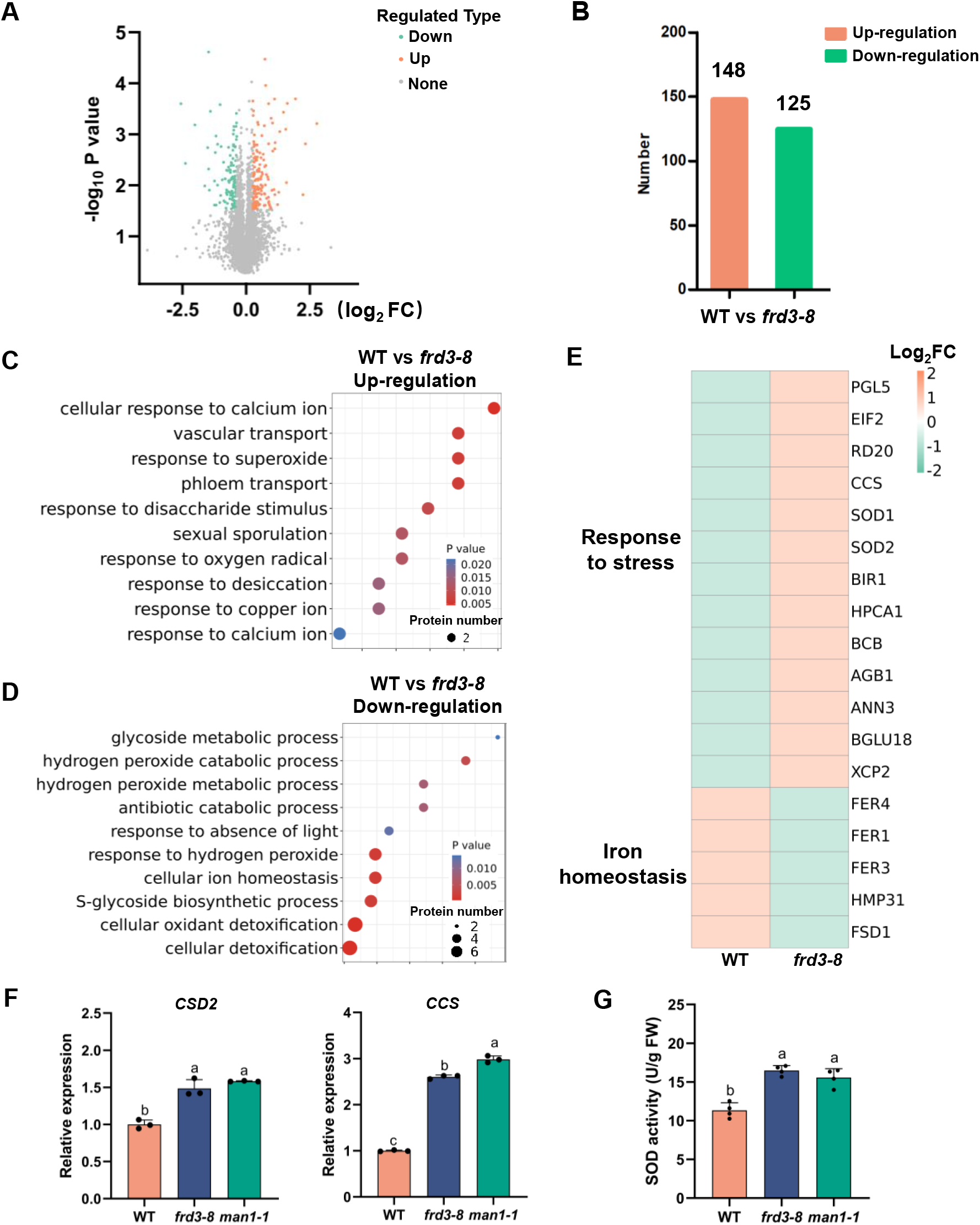
Proteomics analyses *in frd3-8* mutant and wild type. A. Volcano plot of differentially expressed proteins in WT vs *frd3-8* group. Orange dots, green dots, and gray dots indicate the up-regulated, down-regulated, and indifference genes, respectively. B. The number of differentially expressed genes (DEGs) in WT vs *frd3-8* group. Differentially expressed proteins were characterized by the threshold (the absolute value of log_2_ (fold change ≥ 1.2, *P* value ≤ 0.05). C-D. GO terms enrichment analysis. Differentially expressed proteins in WT vs *frd3-8* up-regulated group (C), WT vs *frd3-8* down-regulated group (D) were enriched by GO terms based on biological processes. *P* value ≤ 0.05. E. Hierarchical clustering analysis of the enriched proteins related to iron homeostasis and stress in WT and *frd3-8* lines. F. The transcript levels of *CCS* and *CSD2* genes were measured in 3-week-old soil-grown WT, *frd3-8*, and *man1-1* plants by qRT‒PCR. Values are mean ± SD (n = 3 replicates). Different letters indicate significant differences by one-way ANOVA (*P* < 0.05). G. The SOD activity of the leaves of 3-week-old soil-grown WT, *frd3-8*, and man1-1 plants. Values are mean ± SD (n = 3 replicates). Different letters indicate significant differences by one-way ANOVA (*P* < 0.05).

## Discussion

ROS are known to mediate long-range signaling in plants to adapt to high light, heat, and salt stress (Mittler et al., 2011; Baxter et al., 2014; Gilroy et al., 2014). Rapidly enhanced ROS levels might act as an acclimation signal at the initial phase of stress. In this study, we revealed that loss-of-*FRD3* mutants exhibit enhanced drought resistance with reduced stomatal density and increased ABA content in the leaves (Figure 1, 2A, and 2E), which is associated with higher H_2_O_2_ accumulation in roots and leaves (Figure 3). Grafting experiment results supported that root-derived long-range signals generated in *frd3* rootstock activate ABA synthesis in the leaves by upregulating the expression of *NCED3*, thereby conferring drought tolerance (Figure 4). Furthermore, foliar Fe supplementation of the *frd3* mutant did not revert the drought resistance phenotype to that of the wild type (Figure 5), ruling out the possibility of Fe deficiency causing the drought resistance phenotype. Comparative transcriptomic and proteomic analyses revealed that loss of *FRD3* altered the expression of a set of genes/proteins associated with abiotic stress response, ROS and iron homeostasis, and signal transduction (Figure 7 and 8), consistent with the mutant phenotype.

Our findings suggest that root-derived long-range signals can trigger a drought response cascade in the leaves of *frd3* mutants and that H_2_O_2_ is a strong candidate long-range signal. Our dehydration experimental data imply that H_2_O_2_ may serve as a long-distance signal in the drought response of wild type in general (Figure 6). A working model is proposed to illustrate root-derived long-range H_2_O_2_ signaling in the drought response (Figure 9).

**Figure 9.**
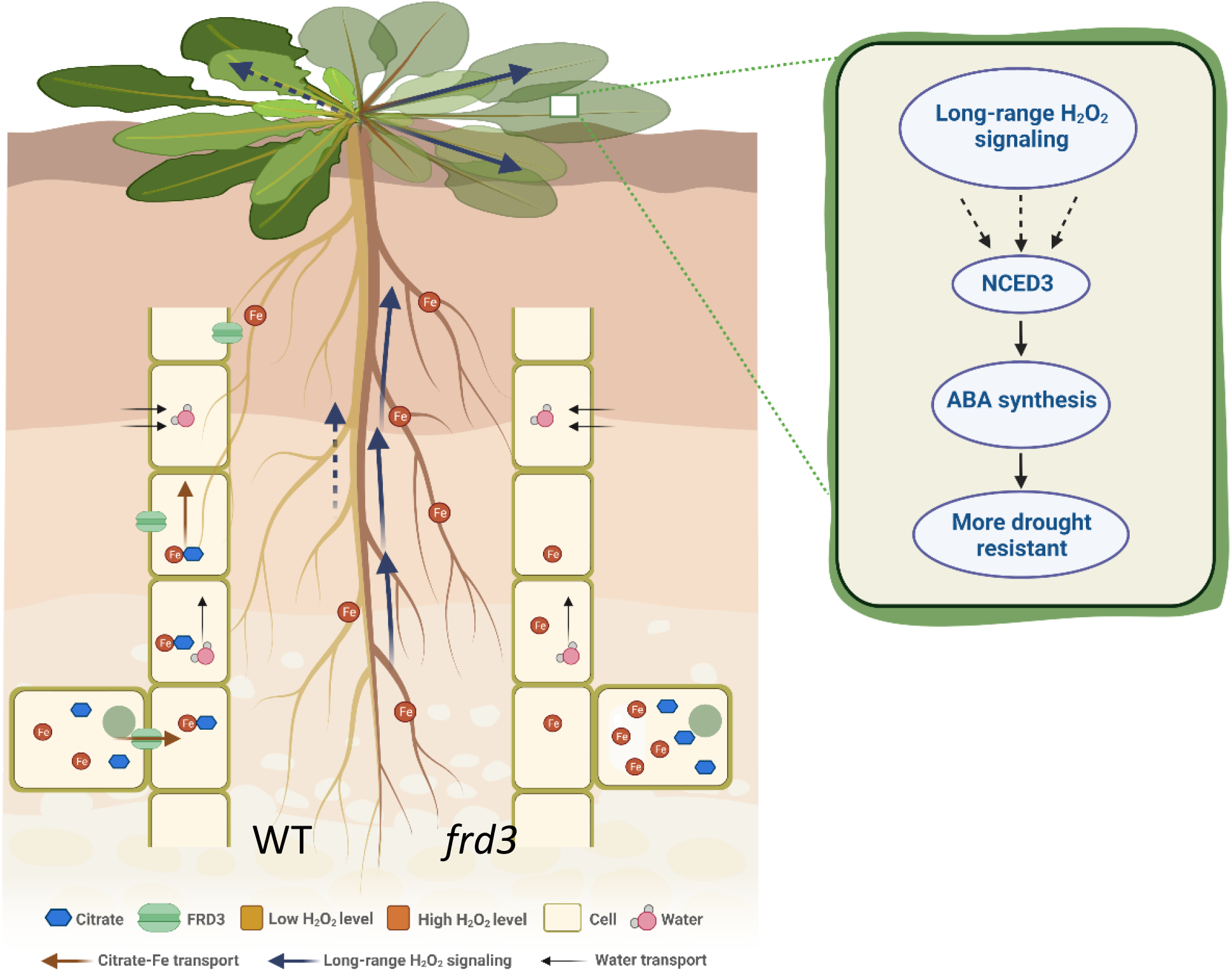
A working model of root-derived long-range signaling confers drought resistance in *frd3* mutants. FRD3 mediates the intracellular efflux of citrate to the apoplast. Loss of *FRD3* caused reduced citrate excretion and excess Fe accumulation in roots. Furthermore, the *frd3* mutants enhanced H_2_O_2_ accumulation in the roots and leaves. Root-derived long-range signaling (H_2_O_2_) activates ABA synthesis by upregulating the transcript of *NCED3* in the leaves of *frd3* mutants with enhanced drought resistance.

### H_2_O_2_ is a strong candidate for long-range signals in *frd3* mutants

Long-distance signaling is important in the drought response of plants, where roots and shoots communicate through hydraulic signals (Christmann et al., 2007) and sulfate (Malcheska et al., 2017; Batool et al., 2018). Salt stress-induced Ca^2+^ waves are also involved in root-to-shoot signaling in plants (Choi et al., 2014). Recently, the CLE25 peptide was reported to function as a root-to-shoot signal to modulate the drought response by affecting ABA biosynthesis and stomatal closure (Takahashi et al., 2018). The ROS-dependent systemic response, an autopropagating ROS systemic signal that travels from the local site to the entire plant, can enhance plant resistance, such as the wounding response (Song et al., 2014). Whether ROS serve as root-to-shoot signals in the drought response is largely uncertain.

Fe is directly involved in ROS production in plant cells (Keunen et al., 2011; Le et al., 2019). Loss of *FRD3* reduces the excretion of citrate, causing excess Fe accumulation in roots (Green and Rogers, 2004; Durrett et al., 2007; Li et al., 2019). In this study, *frd3* mutants showed increased H_2_O_2_ accumulation in roots and leaves (Figure 3A), suggesting that loss of *FRD3* leads to excess Fe accumulation in roots, thus contributing to ROS production, which is supported by the data of transcriptomic and proteomic analyses (Figure 7 and 8). Differentially expressed SOD genes/proteins were upregulated in the *frd3-8* mutant (Figure 7C and 8E). SODs function in scavenging superoxide radicals by catalyzing the dismutation of O_2_^-^ to H_2_O_2_ and O_2_ (Gechev et al., 2006; Gill and Tuteja, 2010). Therefore, elevated SODs in the *frd3-8* mutant may promote H_2_O_2_ production, resulting in an enhancement of H_2_O_2_ signaling.

H_2_O_2_ is known as a key signaling molecule mediating long-range adaptive responses in plants (Mittler et al., 2011; Saxena et al., 2016). NADPH-generated root H_2_O_2_ signals confer salt tolerance and trigger rapid stomatal closure in grafted pumpkin plants (Niu et al., 2018). Our study showed that the grafted plants with *frd3* mutant rootstock caused an increased H_2_O_2_ level in the leaves compared to the wild-type rootstock and improved drought resistance (Figure 4E), indicating that the root-derived signals in the *frd3* mutants trigger drought response, leading to improved drought resistance. Moreover, Fe deficiency, a typical phenotype of the *frd3* mutant, was ruled out as a cause of drought resistance by the foliar Fe supplement that failed to reverse the drought resistance phenotype of the *frd3-8* mutant to the wild type (Figure 5), reconfirming the probability of root-derived H_2_O_2_ signals triggering drought resistance. Additionally, dehydration triggered an H_2_O_2_ increase in the roots and leaves of the wild type (Figure 6). Together, these results suggest that H_2_O_2_ may act as a long-range signal not only in the *frd3* mutant but also in the wild type.

### How do root-derived long-range signals trigger ABA synthesis in the leaves of *frd3* mutants?

ROS mediate rapid cell-to-cell communication over long distances under abiotic stress, which plays a vital role in regulating rapid SAA (Miller et al., 2009; Mittler and Blumwald, 2015). However, how the root-to-shoot ROS signal is perceived by the leaves to trigger the drought response is still unknown. Glutathione peroxidase-like enzyme (GPX3) serves as a sensor and transducer of H_2_O_2_ signaling, which mediates ABA and oxidative signaling to regulate the drought stress response (Delaunay et al., 2002; Miao et al., 2006). Recently, receptor-like kinases have been considered candidate ROS sensors because they can sense ROS through redox modifications of the extracellular structural domain (Tripathy and Oelmüller, 2012). HYDROGEN PEROXIDE INDUCED CALCIUM INCREASES 1 (HPCA1), a leucine-rich repeat receptor kinase acting as a hydrogen peroxide sensor, was shown to play a key role in H_2_O_2_-induced activation of Ca^2+^ channels to modulate stomatal closure (Wu et al., 2020). Proteomics analysis showed that HPCA expression was significantly elevated in *frd3* mutants (Figure 8E), suggesting that *frd3* root-derived long-distance signals possibly depend on HPCA to activate downstream events. Moreover, receptor-like kinases localized on the plasma membrane, GHR1 (Hua et al., 2012) and RDK1 (Kumar et al., 2017), which modulate drought resistance in an ABA-dependent manner, were upregulated in the plants with *frd3* mutant rootstock (Figure 7C). It was previously demonstrated that H_2_O_2_ functions as a potent activator of MAPK cascades, playing key roles in the induction of the stress response (Kovtun et al., 2000; Mittler et al., 2022). In this study, we showed that the transcript abundance of *MAPKKK14* in *frd3-8*/*frd3-8* and WT/*frd3-8* plants was higher than that in WT/WT plants. Therefore, we speculate that these receptor-like kinases may act as H_2_O_2_ sensors and affect ROS-related signaling cascades.

A previous study showed that ABA accumulation in systemic acclimation to heat stress is RBOHD dependent, indicating that ABA is modulated by the systemic ROS signal (Suzuki et al., 2013). Furthermore, ROS are thought to promote ABA biosynthesis, while increased ABA levels also lead to ROS production in guard cells, which forms a positive feedback loop between ROS and ABA for rapidly modulating stomatal closure (Song et al., 2014; Mittler and Blumwald, 2015). Our grafting experiments showed that the higher H_2_O_2_ levels in the *frd3* rootstock trigger H_2_O_2_ accumulation in the scion leaves of the wild type through long-distance translocation (Figure 4). H_2_O_2_ may somehow modify and activate key components in the ABA signaling pathway, which upregulates the expression of *NCED3* in the leaves and increases ABA levels (Figure 4). Transcriptomic analyses also revealed that *NCED3* and *ABA2* were upregulated in *frd3-8*/*frd3-8* and WT/*frd3-8* grafted plants (Figure 7C). The CLE25 peptide has been reported to move from roots to shoots to enhance ABA accumulation by inducing *NCED3* expression in the leaves (Takahashi et al., 2018; Takahashi et al., 2020). Drought stress induces the expression of *NCED3* in *cle25*/WT grafted leaves but not in WT/*cle25* grafted plants. Nevertheless, the presence of retrograde signaling pathways caused by ROS and ROS-induced oxidative modifications of target proteins show that ROS can deliver and affect gene expression in the nucleus and ultimately affect the entire plant (Pornsiriwong et al., 2017; Li and Kim, 2021). Whether the expression of *NCED3* for ABA biosynthesis in *frd3* mutants was activated by ROS or ROS-related signaling cascade components awaits further investigation. Furthermore, dehydration triggered H_2_O_2_ accumulation in the roots and leaves of the wild type (Figure 6), implying that H_2_O_2_ may serve as a long-range dehydration signal in the drought response.

### Persistent high ABA reduces stomatal density

ABA is well known to control transpiration loss under drought stress by regulating stomatal movement. However, increasing evidence suggests that ABA may also regulate stomatal density. It is well known that ABA limits the initiation of stomatal lineage development in *Arabidopsis* leaves by negatively regulating *SPEECHLESS* (*SPCH*) and *MUTE* expression (Tanaka et al., 2013). ABA_-_mediated regulation of stomatal density is OST1_-_independent (Jalakas et al., 2018). Recently, the core SnRK2 kinases (SnRK2.2, SnRK2.3, SnRK2.6) of ABA signaling were reported to directly phosphorylate SPCH to suppress stomatal production (Yang et al., 2022). In our study, we showed that loss of *FRD3* causes higher ABA content with reduced stomatal density in the leaves (Figure 2), suggesting that a persistently higher leaf ABA content may reduce stomatal density. Meanwhile, our RNA-seq analysis revealed that *ERECTA* (*ER*), encoding LRR receptor-like kinases (LRR-RLKs) that function upstream of the MAPK cascades and control stomatal patterning (Shpak et al., 2005; Qi et al., 2018) and stomatal density (Guo et al., 2019), is significantly upregulated in *frd3-8*/*frd3-8* and WT/*frd3-8* grafted plants (Figure 7C). Environmental signals likely converge with the MAPK module to influence stomatal density and distribution (Pillitteri and Torii, 2012). Thus, we speculate that the higher ABA levels in the *frd3* mutant would enhance ABA signaling and affect *ER* expression and MAPK cascades, resulting in reduced stomatal density.

## Materials and Methods

### Plant materials and growth conditions

All *Arabidopsis thaliana* plants used in this study were the Columbia-0 ecotype (WT). The *frd3-8* mutant (SALK_122235C) and *man1-1* mutant (CS8506) were obtained from the Arabidopsis Biological Resource Centre (ABRC). The homozygous *frd3-8* mutant was verified by RT PCR analysis with the specific primers listed in Table S1. The *man1-1* mutant was confirmed by sequencing. To generate the overexpression lines (OX-3, OX-4), the construct *35Spro::FRD3* was made by inserting *FRD3* cDNA into the pCB2004 vector, which was used to transform *Arabidopsis* plants using the floral-dip method (Clough and Bent, 1998).

The seeds were sterilized with 10% bleach for 15 min and then washed four times with sterile distilled water. Seeds were treated for 2 days at 4°C in the dark and then vertically germinated on MS (Murashige and Skoog) medium with 1% (w/v) sucrose at 22°C and 60%∼80% relative humidity with a 16-hour light/8-hour dark photoperiod.

### Grafting experiments

The grafting assay was performed as previously described (Li et al., 2019). The cotyledons were removed from the seedlings vertically grown in MS medium for 6 days, and then hypocotyls were cut from 1/3 of the upper part and divided into two parts: rootstock and scion. The corresponding rootstock and scion were gently combined on new MS medium. After horizontal growth for 10 days, the seedlings without adventitious roots were transferred to the soil for further cultivation and used for subsequent experiments.

### RNA isolation and quantitative RTDPCR (qRTDPCR)

Total RNA was extracted using *TransZol* reagent (TransGen Biotech, China), and then reverse reaction was performed using TransScript® II First-Strand cDNA Synthesis SuperMix (TransGen Biotech, China). For reverse transcription-polymerase chain reaction (RT PCR) analysis, first-strand cDNA was synthesized from 1 µg of mRNA, and then the cDNA was diluted to a certain concentration as a template for RT PCR amplification and quantitative RT PCR. *UBIQUITIN5* (*UBQ5*) was used as an internal control, and the experiment was performed in three biological replicates, each with three technical replicates. The primers used in the experiment are shown in Table S1.

### Histochemical GUS staining

To generate the *FRD3pro::GUS* reporter line, the construct was made by inserting the 2 kb promoter of *FRD3* into the pCB308R vector, which was used to transform *Arabidopsis* using the floral-dip method.

Different tissues of *FRD3pro::GUS* reporter line plants were incubated in GUS staining solution at 37 in darkness, decolorized with 30%, 50%, 70%, and 100% ethanol for 15 minutes, and rehydrated with 70%, 50%, and 30% ethanol for 15 minutes. Samples were stored in 30% ethanol for subsequent observation.

### Drought resistance assay in soil

Seedlings grown vertically for 6 days on the medium were transferred to soil for growth under 12 hours-light/12 hours-darkness. The seedlings were irrigated with water in the first week and then subjected to drought stress treatment by withholding irrigation. Plants with sufficient watering were used as controls.

For soil water loss, each pot was weighed (m_i_). The subsequent growth conditions were consistent with the drought analysis, and individual pots were weighed again 15 days after the drought treatment (m_15d_). Soil water loss = (m_i_ - m_15d_)/m_i_ × 100%.

### Dehydration assay

Wild-type seedlings grown in MS medium for 7 days were exposed to air drying for 0, 5, 10, and 15 min, whereas the seedlings receiving the control treatment remained in the MS medium. At the end of the treatment, all plants were subjected to DAB staining and observation under the same conditions.

### Stomatal density and sensitivity analysis

The abaxial epidermis was obtained from the 5^th^ and 6^th^ leaves of 3-week-old seedlings as described (Zhang et al., 2004), incubated in an opening solution (10 mM MES-K^+^, 50 mM KCl, 100 μM CaCl_2_) under a light intensity of 100 mol μm^−2^ s^−1^ for 3-4 hours, and then transferred to the closing solution (5 mM MES-K^+^, 10 mM KCl, M CaCl_2_) with 0 or 5 µM ABA for 2 hours. Images were taken with a HiROX MX5040RZ microscope and used for statistical analysis.

### ABA measurement

Leaf ABA quantification was conducted using 3-week-old plants (≥ 0.5 g) according to a previous method (Yang et al., 2001) with enzyme-linked immunosorbent assay (ELISA) kits. The absorbance was detected as described (Yu et al., 2017).

### DAB and NBT staining

DAB and NBT staining for H_2_O_2_ and superoxide in plant tissues were conducted as previously described (Schraudner et al., 1998). Briefly, 7-day-old seedlings or 3- week-old detached rosette leaves were incubated in DAB/NBT phosphate buffer at room temperature in darkness for three hours or twelve hours and then destained with a bleach solution (ethanol: acetic acid: glycerol = 3:1:1). Samples were imaged with a HiROX MX5040RZ microscope.

### Foliar Fe supplement assay

Seven-day-old wild-type and *frd3-8* plants grown in soil side by side were sprayed with 0.15 mL Fe(II)-EDTA solution (0.05 mM; 0.5 mM in 0.01% Silwet L-77 solution) per seedling every 36 h, 7 times before being subjected to the drought assay, while H_2_O (containing 0.01% Silwet L-77) was sprayed as a control treatment.

### Determination of chlorophyll content

The chlorophyll content was determined using a spectrophotometric method. Approximately 0.5 g of leaves were ground and then rinsed and fixed with 3 mL 80% (V/V) ethanol. Subsequently, the supernatant was removed after 3 h of dark extraction to measure the absorbance values at 663 nm and 645 nm. The total chlorophyll content was calculated as Vt is the extraction volume (mL), and FW (g) is the fresh weight. C(mg/g) = (20.21 × A654 + 8.02 × A663) × Vt/FW/1000

### RNA-seq analyses

Sequencing was performed using *Arabidopsis* leaves that had been transferred to soil for three weeks after grafting. Twenty grafted plants of each combination were taken as one sample, and three independent biological replicates were sampled. The RNA-seq library was sequenced by Beijing Biomarker Technology Company (Beijing, China). For the RNA-seq data, differentially expressed genes (DEGs) were characterized by the threshold (the absolute value of log_2_ (fold change) ≥1.5, FDR ≤ 0.01). Gene Ontology (GO) term enrichment analysis for each DEG was conducted using DAVID tools (https://david.ncifcrf.gov/).

### Proteomic analyses

*Arabidopsis* leaves from 3-week-old *frd3* mutant and WT lines were used for proteomics analyses by PTM BIOLABS (Hangzhou, China) using 4D-Label Free technology. Differentially expressed proteins were characterized by the threshold (the absolute value of log_2_ (fold change ≥ 1.2, *P* value ≤ 0.05).

### SOD activity analysis

SOD activity analysis was performed according to a previous method (Peskin and Winterbourn, 2017). Three-week_-_old seedling leaves were used to determine SOD activity, which was measured with a kit and the manufacturer’s instructions (Beijing Solarbio Technology, China).

## Supplemental data

Figure S1. Verification of the *frd3* mutant and *FRD3* overexpression lines.

Figure S2. *FRD3* is preferentially expressed in the hypocotyl and the vascular bundle of the roots.

Figure S3. KEGG pathway analysis of DEGs in the different pairwise comparison groups.

Figure S4. The classification of subcellular localization.

Table S1. Primers were used in this study.

## Author contributions

Q.L. and C.X. designed the experiments. Q.L. performed most experiments and data analyses. P.Z., J.X., and J.W. participated in some experiments and data analyses.

Q.L. wrote the manuscript. P.Z. and C.X. revised the manuscript and supervised the project.

## Acknowledgments

This study was supported by grants from the National Natural Science Foundation of China (31900230 to P.Z. and the China Postdoctoral Science Foundation (2020T130634 and 2019M652200 to P.Z.). We thank the ABRC for *FRD3* mutant seeds (*frd3-8*, *man1-1*).

