## Supplemental info for "Root-derived long-range signals activate ABA synthesis in *frd3* leaves to enhance drought resistance"

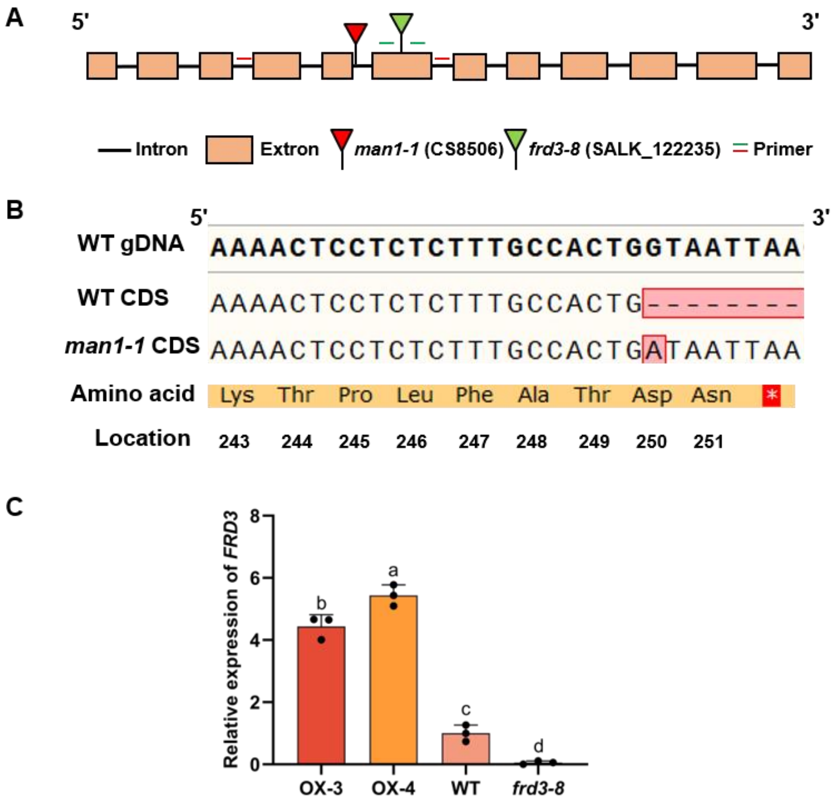


**Figure S1. Verification of the *frd3* mutant and *FRD3* overexpression lines**

A. Schematic illustration of the *frd3-8* mutant and *man1-1* mutant used in this study. The *frd3-8* is a mutant in which a T-DNA insertion into the sixth exon causes premature termination of transcription. The *man1-1* mutant was produced by EMS mutagenesis.

B. Sequence confirmation of *man1-1* mutant. The *man1-1* mutant is homozygous with a G-to-A transition in the first nucleotide of the fifth intron resulting in the retention of the intron and shifting of the reading frame, leading to premature termination with a stop codon.

C. *FRD3* transcript levels of *35S:FRD3* lines (OX-3 and OX-4) and the *frd3-8* mutant. Seeds of WT and 35S:*FRD3* overexpression lines and *frd3* mutants germinated and grown on MS medium for 7 days, then RNA was isolated, and quantitative RT−PCR analyses were performed to detect the transcript level of *FRD3*. Values are mean ± SD (n = 3 replicates). Different letters indicate significant differences by one-way ANOVA ($P < 0.05$).

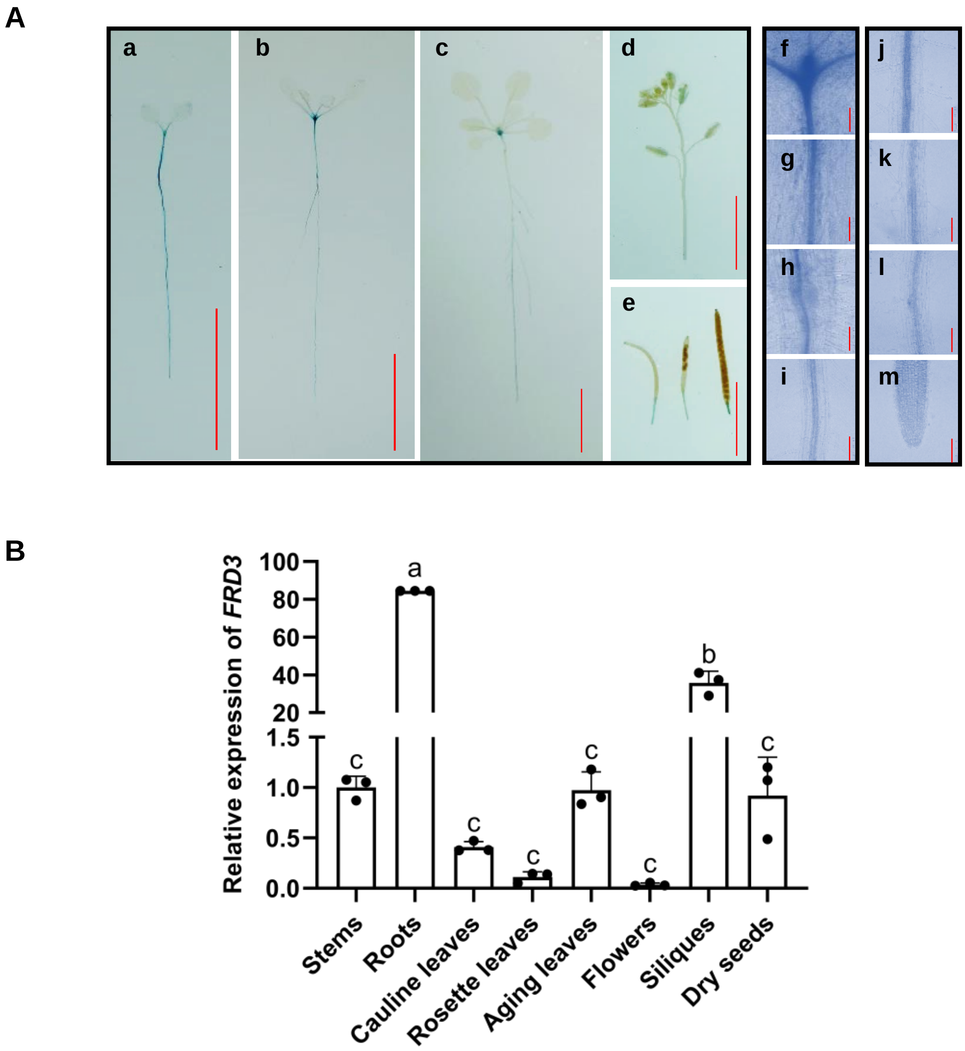


**Figure S2. *FRD3* is preferentially expressed in the hypocotyl and the vascular bundle of the roots**

A. The GUS staining of *FRD3pro:: GUS* transgenic plants. The GUS activity was observed in the seedings with two true leaves (a), two cotyledons (b), six cotyledons (c), flowers (d) and siliques (e), and the rhizomes junction (f), hypocotyl (g), and primary root (h-m). These different tissues were incubated in the GUS staining solution for 2 hours before photographs were taken (A). Bar = 1 cm in a-e, bar = 100 μm in f-m. At least 30 independent lines were used for the GUS staining.

B. *FRD3* transcript levels in different tissues. Different tissues of 6-week-old bolting WT plants were used to isolate RNA, and the transcript level of *FRD3* was detected by quantitative RT−PCR analysis. Values are mean ± SD (n = 3 replicates).

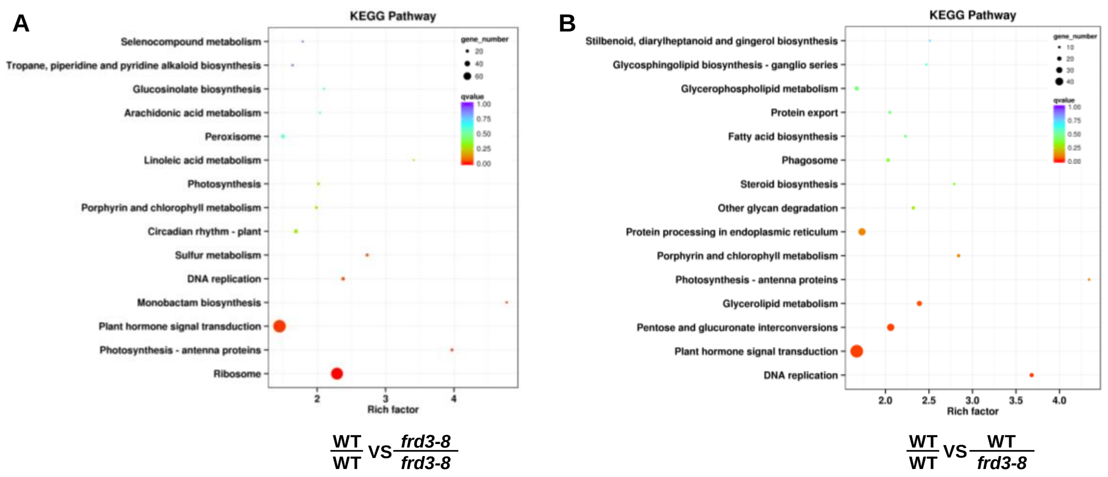


**Figure S3. KEGG pathway analysis of DEGs in the different pairwise comparison groups.**

A-B. KEGG pathway analysis of the DEGs (WT/WT vs *frd3-8/frd3-8*, A) and (WT/WT vs WT/*frd3-8*, B). The comparative results of both groups showed that the plant hormone signal transduction was enriched for the most genes with the most significant differences. The abscissa is the Rich factor, which represents the ratio of the proportion of differential genes annotated to a pathway to the proportion of all genes annotated to that pathway, and the ordinate is each KEGG metabolic pathway. The color of the column represents the qvalue of the hypergeometric test.

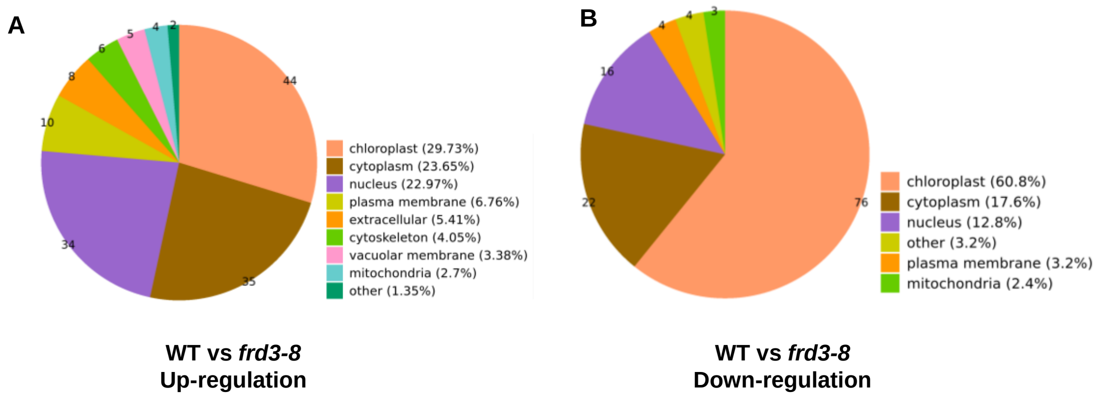


**Figure S4. The classification of subcellular localization**

A-B. The classification of subcellular localization in WT vs *frd3-8* up-regulated group (A), WT vs *frd3-8* down-regulated group (B).

**Table S1. Primers were used in this study.**

| Name | Sequence (5’ to 3’) |
|---|---|
| SALK_122235-LP | ATTTGCAGTTTGGAAGGTTCC |
| SALK_122235-RP | TGCATACCATGGAGTATGCAG |
| CS8506-LP | GACTAATTTGAGTTGAATTTGGATAAC |
| CS8506-RP | TCCTACGACGGATCTTGTTT |
| LBb1.3 | ATTTTGCCGATTTCGGAAC |
| FRD3-qPCR-LP | AGCCACAGCAACCTCCAGCT |
| FRD3-qPCR-RP | TCACTCCCATGACGCCTAGA |
| UBQ-qPCR-LP | AGAAGATCAAGCACAAGCAT |
| UBQ-qPCR-RP | CAGATCAAGCTTCAACTCCT |
| NCED3-qPCR-LP | ATGGCTTCTTTCACGGCAACG |
| NCED3-qPCR-RP | GCGGGAGAGTTTGATGATTGC |
| CCS-qPCR-LP | GAGCCATGCCTCAGCTTCTTAC |
| CCS-qPCR-RP | GGACAAATCCACCTCCACTTTCTC |
| SOD2-qPCR-LP | CAGGGCCTCATGGATTTCATCTCC |
| SOD2-qPCR-RP | TGGAGCTCCGTGTGTCATGTTG |
| COR15B-qPCR-LP | AAGAGTGAGCTCGTCGTCGTTG |
| COR15B-qPCR-RP | TCAGAAGCTTTCTTTGTGGCTTCG |
| ABR1-qPCR-LP | GCTTCAACCGAAGCACAACCTG |
| ABR1-qPCR-RP | TGAACCCGAGTTCCTTGACTGG |
| HAI1-qPCR-LP | ATGGCCATGGCTGTTCCCATGTAG |
| HAI1-qPCR-RP | TCCCAGTCAGCATCAGCTTCAAAC |
| GLR3.6-qPCR-LP | AGAGAGACTACCACCAGCAACC |
| GLR3.6-qPCR-RP | GCTGATGAACTGTGAGGATTGACG |
